# Predicting the operon structure of the *Mycococcus xanthus* genome using the novel software DiscOperon

**DOI:** 10.64898/2026.08.20.745974

**Authors:** Theo Brunet, Bianca H. Habermann

## Abstract

**Motivation:** *Myxococcus xanthus* is a predatory soil bacterium with a large genome of 9.14 MB due to a genome duplication event. While complete genome sequences of *M. xanthus* are available, gene annotation remains challenging due to its size and the resulting large number of duplicated genes. Operons, so syntenic block of genes that are co-regulated in bacterial genomes, are an important resource to help predict gene function accurately.

**Results:** In order to help improve the annotation of complex genomes such as the one from *M. xanthus*, we developed a novel operon prediction tool, DiscOperon, which combines gene expression data with homology searches to identify syntenic blocks: Co-expression data of neighbouring genes across the genome are first used to define gene clusters, which are then used to search for conserved syntenic blocks in fully sequenced bacterial genomes using sequence homology searches. This strategy enables DiscOperon to account for gene insertions, rearrangements and deletions, which is its most distinguishing feature. We have tested DiscOperon against ground truths gene pair information on 3 different species from ODB and RegulonDB and compared it to state-of-the-art and still available operon prediction software and we demonstrate its general usability for operon prediction of any bacterial complete genome. We have applied DiscOperon to predict the operons of *M. xanthus*, which we are making available for the research community.

**Availability and implementation:** DiscOperon is lightweight, user-friendly python tool with minimal dependencies. It is freely available at https://gitlab.com/habermann_lab/discoperon for general usage.

**Contact:** Theo Brunet; Bianca Habermann.

**Supplementary information:** The operon-structured and annotated *M. xanthus* genome is available from this manuscript, as well as from Zenodo (https://doi.org/10.5281/zenodo.21976167). We furthermore plan to submit the *M. xanthus* operon information to the operon database OBD.

## INTRODUCTION

*Myxococcus xanthus* (*M. xanthus*) is an important predatory bacterium mostly found in soil. It can feed on a multitude of other bacteria, as well as fungi. It is a social bacterium and undergoes several lifecycle stages, which include vegetative growth, to fruiting body formation and sporulation; it is capable of explorative as well as social swarming, kin recognition or predation in a group (for a very recent and comprehensive review see (Contreras-Moreno *et al*. 2024)). One of its remarkable features is its wolf-pack-like hunting behaviour and the typical rippling patterns the colony creates while feeding on prey (Thiery and Kaimer 2020). The genome of *M. xanthus* is complex, partially due to the fact that it has undergone a genome duplication event, resulting in a large genome for a bacterium, with 9.14 Mbp and 7,408 protein-coding genes (Jain, Habermann, and Mignot 2021). The annotation of its genes and their correct functional assignments is challenging. Knowing the operon structure of its genome and thus, the association of its genes, can help to obtain a high-quality genome annotation for this organism.

Operons are fundamental structures of regulation in bacterial and archaeal genomes. The first operon described was the Lac operon in *Escherichia coli* through the Lac operon (Jacob *et al*. 1960, Zubay and Lederman 1969). An operon consists of a cluster of adjacent genes transcribed as a single polycistronic mRNA under the control of one promoter. This principle of combining a spatial neighbourhood with transcriptional regulation allows bacteria to coordinate the expression of functionally related genes, often encoding enzymes in the same metabolic pathway or proteins that form protein complexes (Dandekar 1998, Bervoets and Charlier 2019, Zinani, Keseroğlu, and Özbudak 2022, Shine *et al*. 2024). Moreover, by coupling the transcription of multiple genes, operons provide an efficient mechanism for rapid responses to environmental changes (Liang *et al*. 2021).

Operons are also important structures in evolutionary genomics. One well-supported theory based on comparative studies suggests that transcriptional co-regulation is a major driver of operon emergence, especially when the gene regulation has a complex pattern (Price *et al*. 2005). Other theories include emergence through horizontal gene transfer, the nature of ‘selfish operons’, or chromosomal arrangement of genes based on presence in the same protein complex (Lawrence 1999, Ballouz *et al*. 2010, Zhu, Hong, and Wang 2024). Comparative studies have for instance shown that operons can emerge through gene duplication, horizontal gene transfer or insertion (Price, Arkin, and Alm 2006). While operons are considered dynamic, many of them are also highly conserved, such as the **trp** operon involved in the tryptophane biosynthesis, or the **ara** operon, which is involved in arabinose transport and metabolism (Ogden *et al*. 1980, Merino, Jensen, and Yanofsky 2008). Their conservation is due to a selective pressure to maintain the co-regulation of the hosted genes (Price, Arkin, and Alm 2006). Given their importance, the detection of operons in bacterial genomes is essential to understand their biology. To detect operons experimentally is difficult and time-consuming, which is why computational methods are needed to predict them.

Several tools and databases have been created in order to detect operons and to store operon information for different bacteria, which are all based on different operon features (reviewed in (Zaidi and Zhang 2016, a non-comprehensive list of methods can be found in **Supplementary Table S1**). Among these features are predominantly genomic ones, such as transcription start or termination sites, intergenic distances between genes, or directionality of the genes; they also include gene functional features or protein-protein interactions, conserved genomic arrangements across species, as well as gene expression data, or visual representation of bacterial genomic features (e.g. reviewed in (Chuang *et al*. 2012, Zaidi and Zhang 2016)). Many of the developed tools are web-based, such as OperonFinder (Tomar, Dasgupta, and Kanaujia 2023), which uses deep learning for operon prediction in bacteria or archaea, or Operon-mapper (Taboada *et al*. 2018). There exist also web-based resources for operons, such as the ODB database (Okuda and Yoshizawa 2011) or RegulonDB (Salgado *et al*. 2024), both of which are the most consistently used resources for operon information still available to date. While gene expression data can be used as a very strong indicator of existing operon structures in neighbouring genes, it is rarely combined with evolutionary information on conserved syntenic blocks in the genome to identify potential operons. Moreover, many tools focus exclusively on direct gene neighbours and neglect potential gene loss or rearrangements in longer syntenic blocks.

In this work, we wanted to use operon prediction to improve the genome annotation of the *M. xanthus* genome DK162. To this end, we developed a novel open-source tool for operon detection, DiscOperon, which employs gene expression data in combination with evolutionary information to allow prediction of longer operon structures and to account for gene loss or rearrangements. We compared DiscOperon to some of the best operon prediction tools still available to date and achieved equal or better performance. Using DiscOperon on the *M. xanthus* genome, we identified 1218 operons with a mean gene content of 3.29 genes per operon. In order to allow operon annotation with DiscOperon for many users and for any bacterial genome, we ensured user-friendly and light-weight design of DiscOperon, which runs on a standard computer with no dependence on GPU infrastructure or excessive programming skills of the user.

## METHODS

DiscOperon is an open-source pipeline, designed to predict operon structures in bacterial genomes using transcriptomic data and comparative genomics. The workflow integrates expression correlation, sequence homology searches using DIAMOND (Buchfink, Xie, and Huson 2015, Buchfink, Reuter, and Drost 2021), synteny evaluation, and machine learning classification. The workflow of DiscOperon can be seen in **Figure 1**.

**Figure 1:**
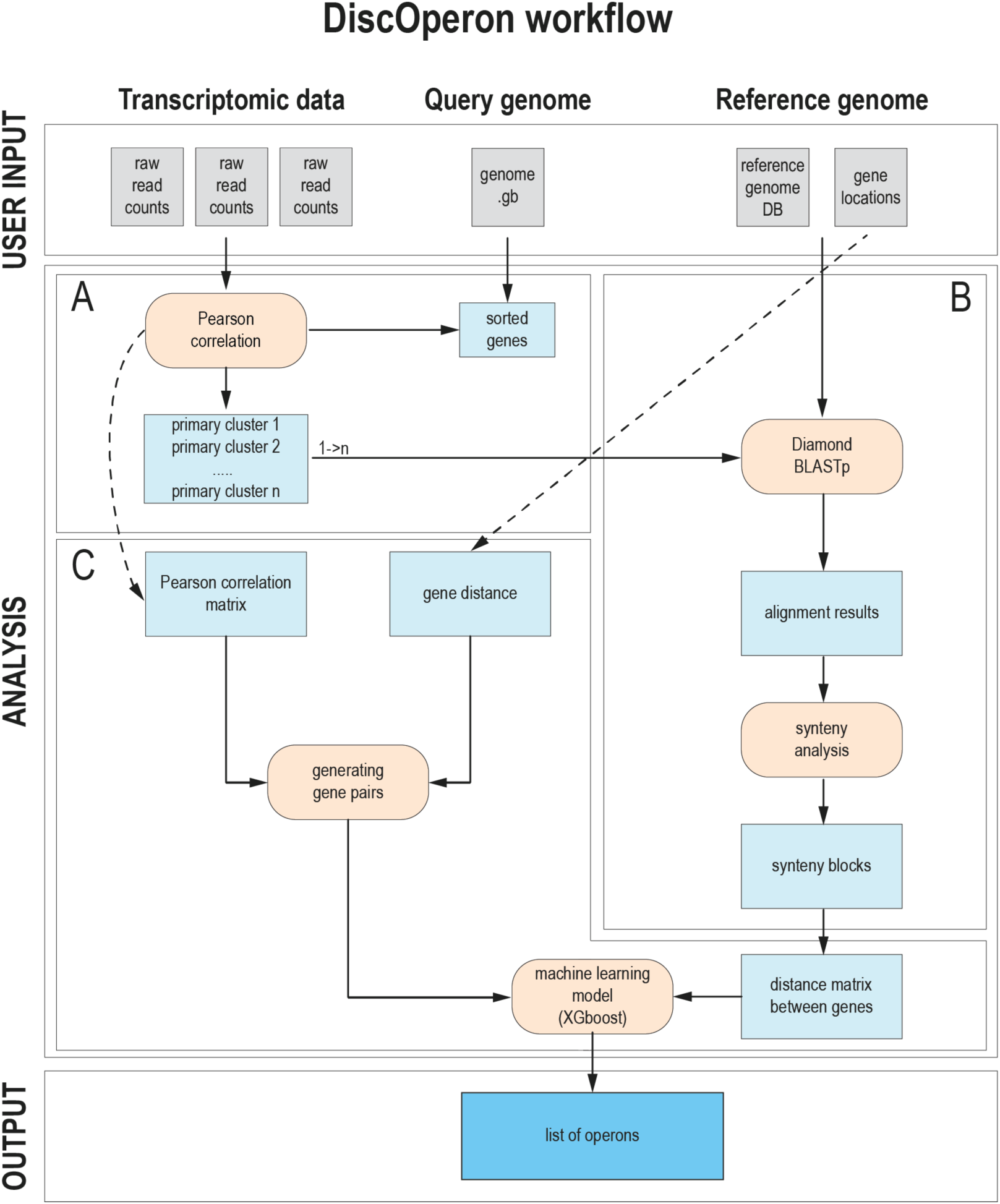
Workflow of DiscOperon. The user needs the query genome of the species of interest, transcriptomic data of the species of interest, as well as reference genomes for comparative genomics analysis as input. **(A)** Transcriptomic data are used to assess co-expression based on pearson correlation and, together with positional information, are used create primary gene clusters. **(B)** Primary clusters are used to perform comparative genomics and synteny analysis using DIAMOND BLASTp. **(C)** The pearson correlation matrix together with gene distances from homologs are used to generate gene pairs, and together with the distance matrix between the genes from synteny analysis is used with an XGboost machine learning model to predict list of operons. The output of the software are the predicted operons with all their genes, so no post-processing is required.

### User-provided input data

DiscOperon needs three kinds of files: 1. a GTF file of the bacterial genome of interest; this file is used to retrieve the genomic coordinates and strandedness of each gene. 2. A fasta-file containing all protein sequences of the bacterium. 3. One or multiple analysed datasets of RNA-seq data that contain at least the gene IDs as the first column and one or multiple feature counts results in the other columns. Both the GTF and fasta file can be easily retrieved from public databases such as ENSEMBL (Dyer *et al*. 2025) or NCBI (Sayers *et al*. 2025). The RNA-seq files can be generated by tools such as featurecounts (Liao, Smyth, and Shi 2014), or, if provided with data submission, also retrieved from public resources such as Gene Expression Omnibus (Clough *et al*. 2024), or ArrayExpress (Athar *et al*. 2019).

### Step 1: Dynamic clustering

The transcriptomic data is first normalized in transcript per million (TPM) and then integrated with the genomic annotation.In order to identify clusters of contiguous genes that are co-expressed, the genes are sorted by genomic location and then split in two groups based on their orientation. Gene co-expression is then computed using the Pearson correlation coefficients on the normalized transcriptomic data and an associated p-value is calculated using the Student t-distribution. To identify operon-like clusters, a dynamic programming algorithm is used to detect contiguous regions that maximize the intra-cluster correlation. This algorithm optimize the scoring function [1]

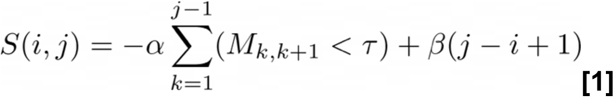

Where S(i,j) is the score of the cluster spanning from the gene of index i to the gene of index j, **ɑ** represent the penalty applied to the pair of genes where the correlation **M_k,k+1_** is lower than the threshold **τ** and **β** is a reward term used to favor larger cluster. This algorithm allows the creation of large clusters of genes that are globally more correlated. Those clusters are then used to facilitate the synteny conservation analysis by doing a first rough slicing of the genome.

### Step 2: Synteny based analysis

Next, homology-based searches are used to identify potential synteny conservation of the different clusters in other bacterial genomes. To this end, we created a blast-database using the *makedb* script from the DIAMOND suite (v 2.1.8.162), using all complete bacterial reference genomes in Refseq. Synteny searches are done using sliced clusters of proteins from the input genome and using the BLASTp like algorithm from DIAMOND, in order to retrieve the homologs of each protein of our query genome in the other bacterial genomes. The BLAST results are filtered based on the sequence length coverage (default: 50 %) and identity (default: 25 %) and only those satisfying both criteria are used for further analysis. Results are sliced based on the dynamic clustering results, producing one file per gene cluster for further processing. Homologous genes identified in each of the genomes are then sorted by genomic position. For each homolog, we retrieve the neighbouring genes that are closer than a defined distance threshold. This step ensures that we know for each of the homologs their genomic context and if multiple homologs are found in the same genomic region. A bit score is produced as output for each cluster, which is the sum of the bit score of each of the homologs that are in the same cluster; potentially neighbouring genes without homologs do not contribute to this score. Finally for each pair of genes in the clusters found in the first step we compute several metrics that are used to determine, if two genes are in the same operon or not:

- their co-occurrence in the other bacterial genomes
- the average base pairs distance between their homologs
- the conservation of the distance between the two genes across the other bacterial genomes
- the maximal base pairs distance between their homologs
- the expression correlation from transcriptomic data
- the genomic rank distance between the two genes
- the number of genes between the two genes
- the genomic span of the two genes in base pairs

### Step 3: Classification of operon content

Finally, we classify the genes, whether, based on the above criteria, they are part of an operon structure, or not. For each pair of genes, DiscOperon retrieves the Pearson correlation and the 7 features from the previous synteny analysis. Those data are then used by a cross-validated calibrated XGBoost model in order to classify each pair as operonic or not. A final reconstruction step is done to create the identified operons by grouping all pairs of genes. The final output of the tool is a file containing a list of predicted operons as well as a score for each pair of genes in an operon.

### Other operon search tools, data and metrics used for method validation

In order to train and test the model we extracted operonic information from two databases, RegulonDB (for *E. coli*) and ODB for *E. coli, Bacillus Subtilis (B. subtilis)* and *Corynebacterium glutamicum (C. glutamicum)*. From the operons in these two databases, we generated a list of pairs of genes that belong to the same operon and that will be used to train and evaluate our new model, DiscOperon, with the following extracted information (**Table 1**):

**Table 1:**
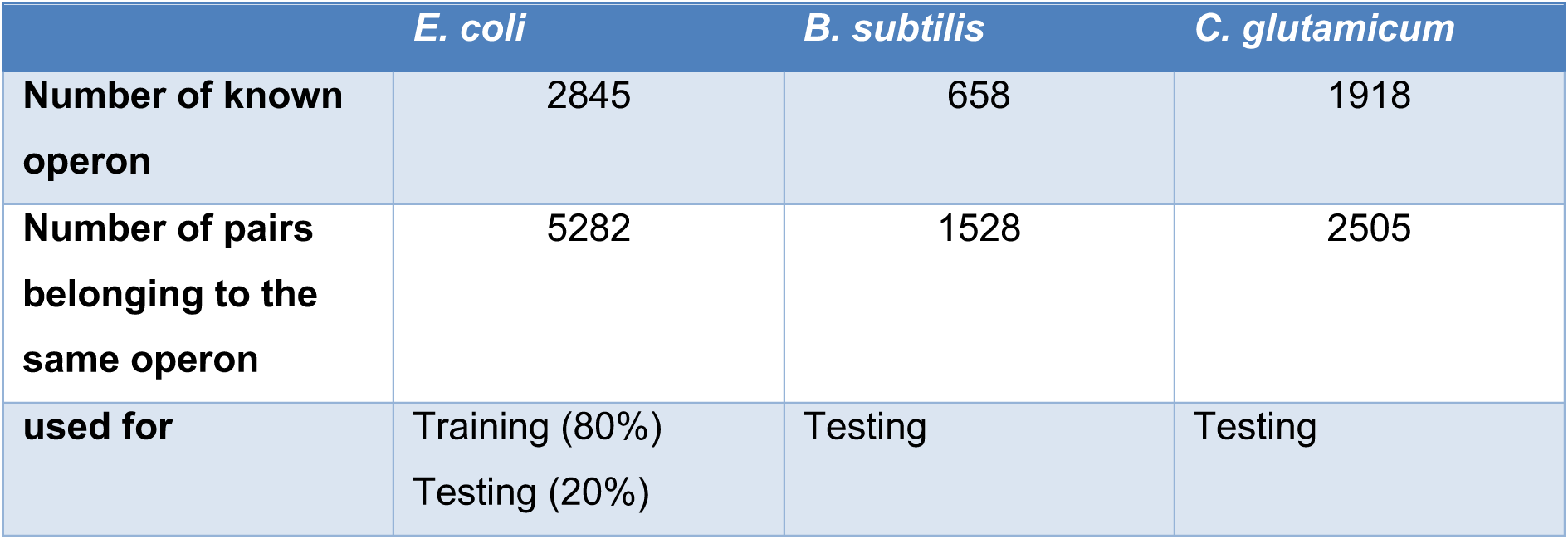
Operon information extracted from OperonDB and RegulonDB used to train and test operon prediction models.

|  | *E. coli* | *B. subtilis* | *C. glutamicum* |
| --- | --- | --- | --- |
| **Number of known operon** | 2845 | 658 | 1918 |
| **Number of pairs belonging to the same operon** | 5282 | 1528 | 2505 |
| **used for** | Training (80%)<br>Testing (20%) | Testing | Testing |

We benchmarked DiscOperon against three established operon prediction tools OpDetect (Karaji and Peña-Castillo 2025), OperonFinder (Tomar, Dasgupta, and Kanaujia 2023) and Rockhopper (Tjaden 2020). All tools were run using their default parameters and were provided with the same input data to ensure a fair comparison, except OperonFinder which does not use transcriptomic data. Following FAIR principles, all files used are publicly available. The genomes and protein sequences were extracted from the Refseq database and RNA-seq data from the sequence read archive (SRA), both from NCBI. The GEO accession numbers of the different RNA-seq experiments can be found in **Supplementary Table S2**.

### Training and testing set

For this study, operon annotations from ODB and RegulonDB were used to construct the training and testing datasets for the three tested species. We extracted all values from the database of pairs of genes belonging to the same annotated operon. To limit class imbalance while accounting for the incomplete DiscOperon annotation, two additional categories of gene pairs were defined: 1. Negative values were generated from pairs of genes that were assigned to different annotated operons but remained within five genes of each other in the genome. Restricting negative pairs to neighbouring genomic regions creates a more challenging classification task and avoids easy classification only relying on the distance between genes. 2. All remaining gene pairs were classified as unknown, either because they were separated by larger genomic distances or because no experimental evidence was available to determine whether they belonged to the same operon, or not.

### Operon conservation and structure analysis

We used NCBI blast+ from the NCBI suite (Camacho *et al*. 2009) to find paralogs for *M. xanthus* genes by blasting the proteins against the *M. xanthus* protein database (strain DK1622, accession number NC_008095.1 downloaded from NCBI). We kept the closest paralog, based on an E-value cut-off of 1e-20, and a score ration of at least 0.5. For comparing operonic arrangements of paralogs, we only considered the closest hit.

fast.genomics searches for the kil system operons were done online at the fast.genomics web-server(Price and Arkin 2024) https://enigma.lbl.gov/fast-genomics-a-fast-comparative-genome-browser-for-diverse-bacteria-and-archaea/), searching against diverse bacteria and archaea. Genes used for fast.genomics searches are indicated in the figure legends.

## RESULT

### Classification performance of DiscOperon across bacterial species

To test the performance of DiscOperon, we used 3 bacterial species with well annotated operons, which we extracted from ODB and RegulonDB (**Table 1**). To evaluate the predictive performance of DiscOperon, a XGBoost classifier was trained using annotated operon pairs from *E. coli*. To train DiscOperon, we used 80% of data from *E. coli* and used the remaining 20% of *E. coli* operonic data, as well as the complete operon datasets of *B. subtilis* and *C. glutamicum* for testing, helping us to assess DiscOperon’s general applicability to other bacterial species. DiscOperon achieved high performance across all evaluated datasets (**Table 2**), reaching a F1-sore of at least 0.86 and a ROC AUC of at least 0.847 (both on the *C. glutamicum* dataset). It performed highest for the *B. subtilis* dataset, where it obtained a ROC AUC of 0.902 (**Figure 2A**, **Table 2**) and an average precision (AP) of 0.965 (**Figure 2B**, **Table 2)**. Using a score threshold of 0.50, DiscOperon achieved a precision of 0.908, a recall of 0.908 and an F1-score of 0.904 for this bacterium. Similar results were observed for *C. glutamicum*, indicating that the classifier remains effective when applied to species that were not included in the training dataset (**Supplementary Figure S1, Table 2**).

**Figure 2:**
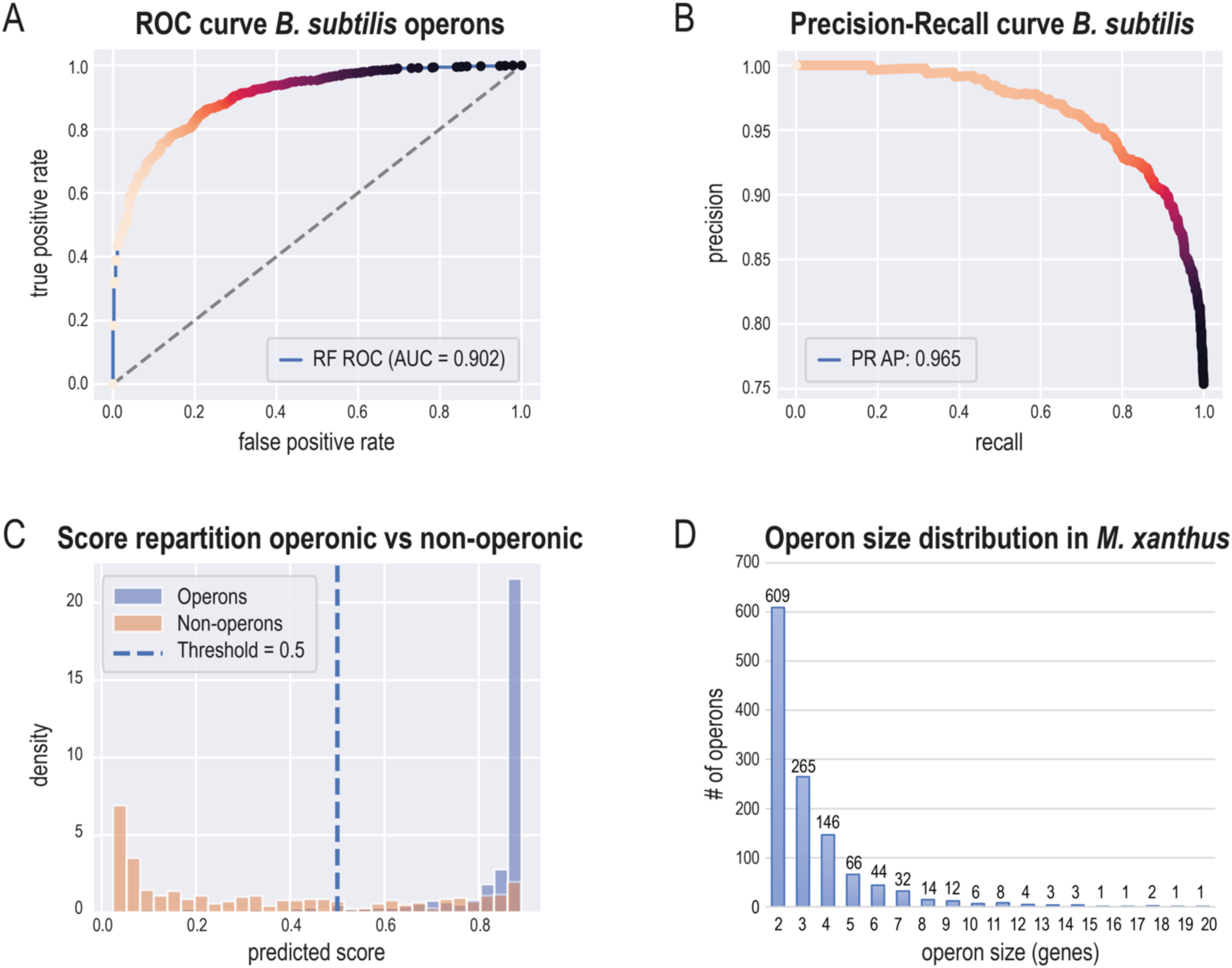
Performance of DiscOperon in correctly predicting operons in *B. subtilis*. **(A)** ROC curve relating true-positive and false-positive prediction rates, and **(B)** Precision-Recall curve of operons predicted for *B. subtilis*. The ROC area under the curve (AUC) was 0.902, the average precision (AP) was 0.965. **(C)** Score repartition of operons vs non-operons. Operons (blue bars) are nearly completely found in a prediction score > 0.5, while non-operons are clearly overrepresented at lower scores, demonstrating DiscOperon’s ability to correctly distinguish operonic vs non-operonic gene pairs. Data for *C. glutamicum* can be found in **Supplementary Figure S1**. **(D)** Gene number distributions of operons detected in *M. xanthus*. Half of operons contain gene pairs, the other half has more than 3 genes. The largest operon has 20 genes and harbours part of the kil system. See also **Supplementary Tables 1-5** for detailed operon information on *M. xanthus*.

**Table 2:**
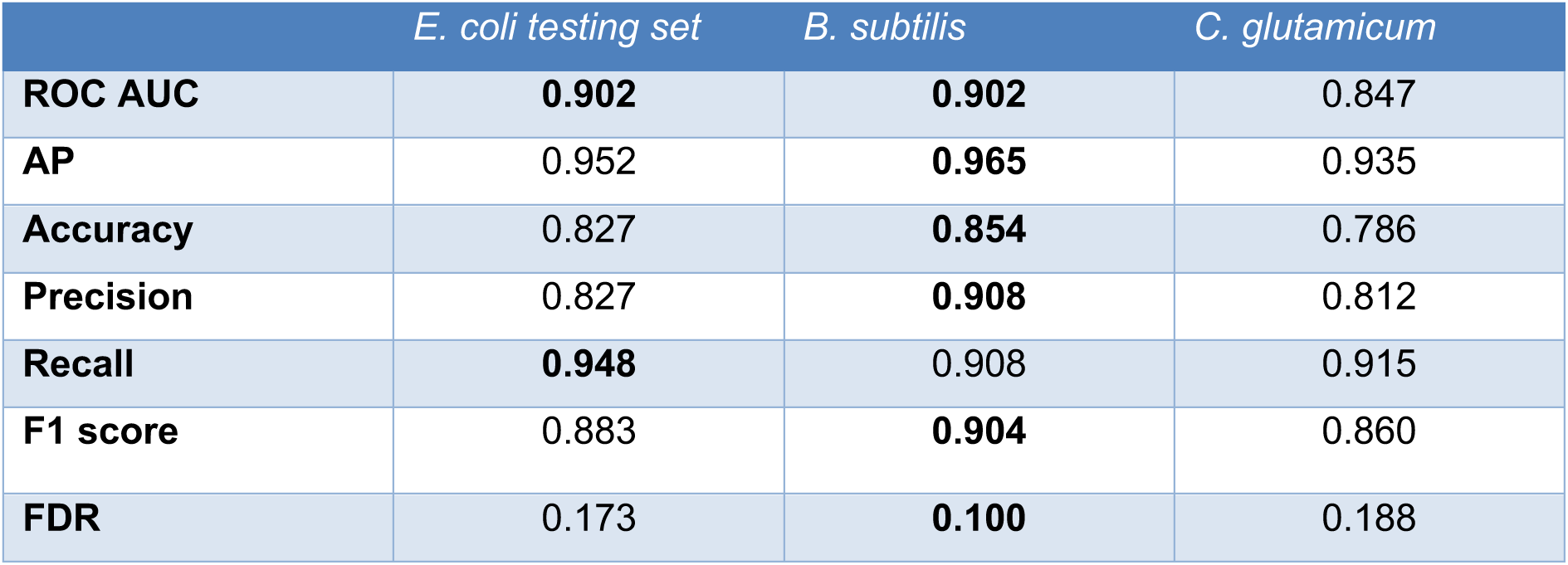
Performance obtained by DiscOperon on predicting operons from different species.

|  | *E. coli* testing set | *B. subtilis* | *C. glutamicum* |
| --- | --- | --- | --- |
| **ROC AUC** | **0.902** | **0.902** | 0.847 |
| **AP** | 0.952 | **0.965** | 0.935 |
| **Accuracy** | 0.827 | **0.854** | 0.786 |
| **Precision** | 0.827 | **0.908** | 0.812 |
| **Recall** | **0.948** | 0.908 | 0.915 |
| **F1 score** | 0.883 | **0.904** | 0.860 |
| **FDR** | 0.173 | **0.100** | 0.188 |

The high precision obtained across species suggests that most predicted operonic gene pairs correspond to true operonic relationships. These results demonstrate that the combination of transcriptomic and synteny-derived features enables accurate operon prediction and can be used on diverse species from different phyla.

### Separation of operonic and non-operonic pairs

To investigate the ability of DiscOperon to distinguish between operonic and non-operonic gene pairs, the distribution of prediction scores was examined, using data from *B. subtilis* (**Figure 2C**). We see a clear separation of operonic pairs at very high scores while the non-operonic pairs seem to be spread at lower scores.

The score density distribution highlights the bimodal nature of the predictions. Most operonic gene pairs accumulated at scores higher than 0.90, while non-operonic pairs were enriched at lower prediction scores. The overlap between the two distributions at low scores indicate that some operonic pairs are harder to classify, either because they are not conserved across species or the expression correlation of the two genes are not strong enough to produce a clear signal.

Based on these score distributions, a threshold of 0.5 was selected for operon reconstruction. This threshold provided a suitable compromise between precision and recall while maintaining a low false discovery rate across the evaluated species. In addition, predictions above this threshold remained highly enriched in known operonic pairs, making it appropriate for the identification of operonic structures.

These results demonstrate that the prediction scores generated by DiscOperon provide a meaningful measure of confidence and can be used to reconstruct operon structures from pairwise classification.

### Comparison against other prediction tools

To benchmark the performance of DiscOperon against existing operon prediction tools, we compared its predictions with those from OperonFinder, OpDetect and Rockhopper, using the same datasets and evaluation metrics (**Table 3**). As these tools only evaluate pairs of adjacent genes, the comparison was restricted to flanking gene pairs. As DiscOperon and several of these tools have been trained on *E. coli,* only *B. subtilis and C. glutamicum* were used for comparison.

**Table 3:**
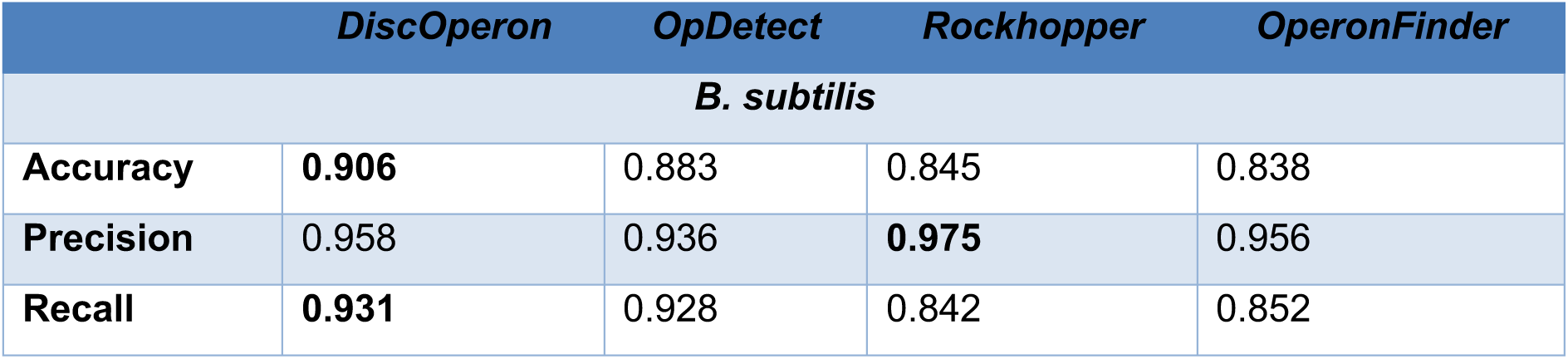
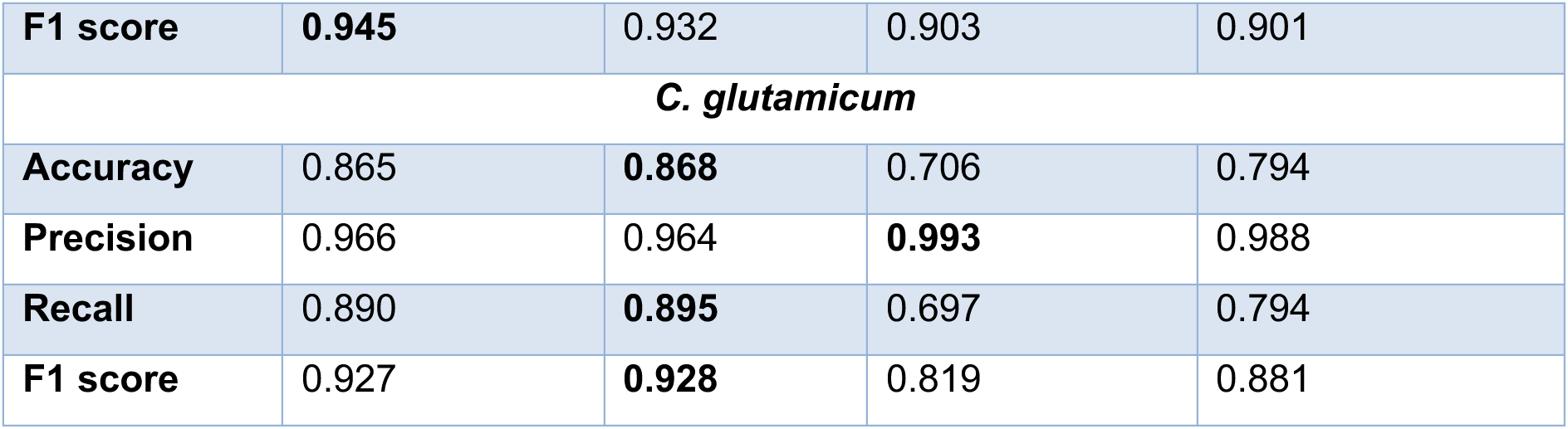
performance comparisons of the different operon prediction methods to predict operonic gene pairs in *B. subtilis* and in *C. glutamicum*.

DiscOperon performed overall better than the other tools across the different metrics, except Rockhopper, which achieved better precision for *B. subtilis* (97.5% vs 95.8%). For *B. subtilis*, DiscOperon performs comparable or better than the other three tools across the different metrics with some exceptions: Rockhopper outperformed DiscOperon in precision (97.5% vs 95.8% for *B. subtilis* and 99.3% vs 96.6% for *C. glutamicum*). For *C. glutamicum*, both OpDetect and DiscOperon achieved similar results, though DiscOperon had a slight decrease in accuracy and recall (0.3% and 0.5%, respectively) compared to OpDetect, as well as a slight increase in precision (0.2%). OperonFinder outperformed DiscOperon for *C. glutamicum* (98.8% vs 96.6%, respectively).

Overall, our results demonstrate that DiscOperon achieves performance comparable or better than the existing operon prediction methods across bacteria from different phyla, validating our approach of combining transcriptomic with evolutionary, as well as syntenic data for operon prediction.

### Operons identified in *M. xanthus*

To further illustrate the ability of DiscOperon to reconstruct operon structure in complete bacterial genomes, we applied our new method to *M. xanthus*. DiscOperon identified 1,218 operons, involving 4,018 genes in total. Predicted operons range from 2 to 20 genes with an average size of 3.29 genes / operon (**Figure 2 D**). There was no extensive conservation of operon structure during genome duplication, at least based on analysis of syntenic arrangements of the closest paralogs (**Suppl. Table S5**). A detailed and annotated list of *M. xanthus* genes and their presence or absence in operons is provided in **Suppl. Table S4**, which could help in improving the functional annotation of genes in *M. xanthus*.

To illustrate the quality of the predicted operons, we examined two well characterised operons involved in different biological processes (the **frz** operon and the **trp** operons), as well as the operons of the **kil** system, which was partially defined previously (Herrou *et al*. 2025).

The **frz** operon encodes the proteins involved in the Frz chemosensory pathway and regulates the frequency of cell reversal (Trudeau, Ward, and Zusman 1996). It consists of two transcriptional units, one containing frzZ and one containing frzA, frzB, frzCD, frzE, frzF and frzG. DiscOperon successfully predicted the two transcription units, frzZ alone and a transcriptional unit containing the 6 other genes **(Figure 3A)**.

**Figure 3:**
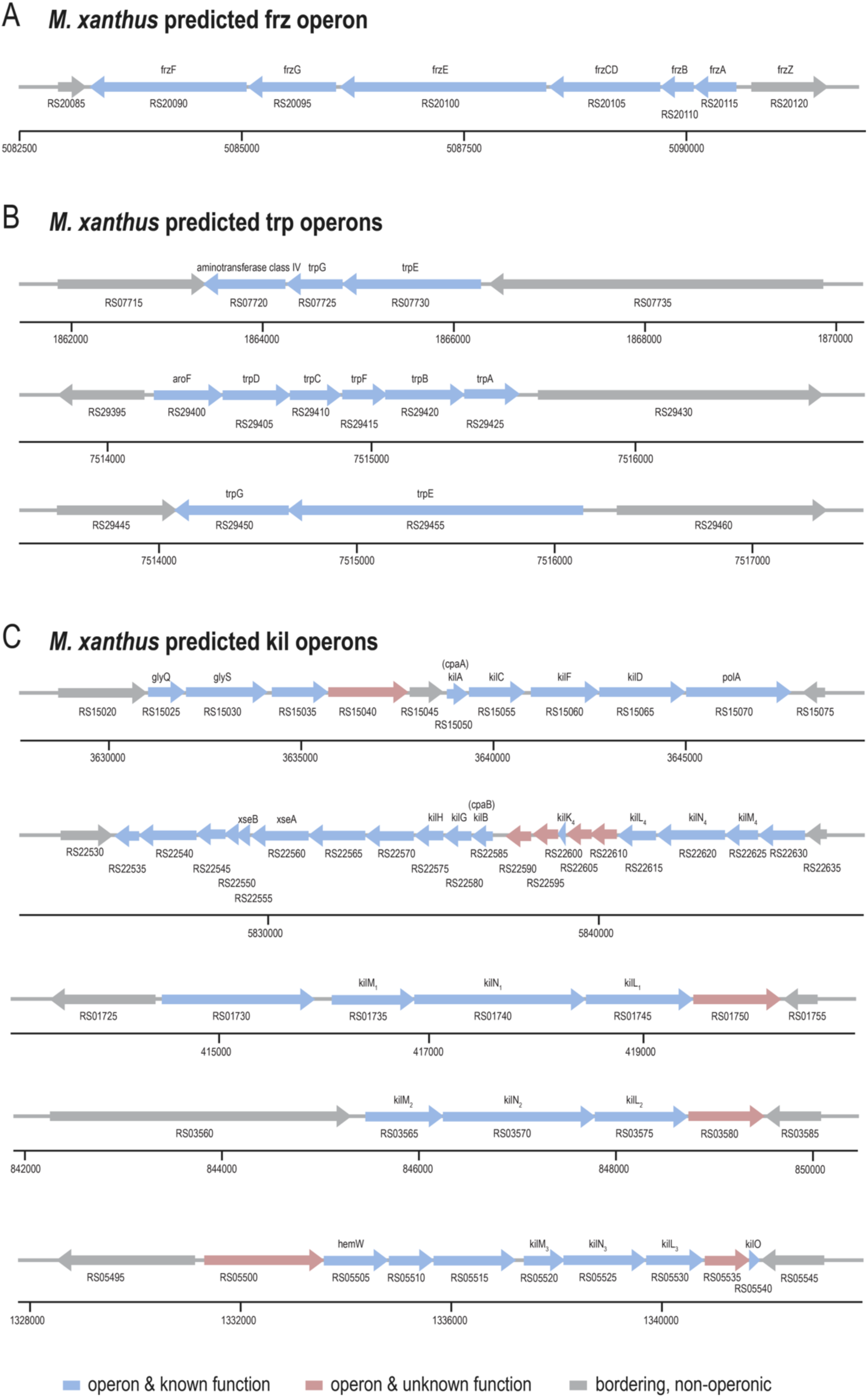
Examples of operons predicted in *M. xanthus*. **(A)** All 6 genes required for the chemosensory Frz pathway are encoded in a single operon in *M. xanthus*, the predicted frz operon. **(B)** Genes involved in the *M. xanthus* Tryptophane biosynthesis pathway are split in 3 different operons. The largest trp operon contains next to aroF, the genes trpD, trpC, trpF, trpB and trpA. The trpE and trpG genes are duplicate in the genome and located in two operons, whereby one with genes MXAN_RS29450 and RS29455 is in close proximity of the main operon and the other contains MXAN_RS07725 and MXAN_RS07730 are predicted to have in addition a class IV aminotransferase, of which only one version exists in the *M. xanthus* genome. **(C)** Operons of the *M. xanthus* kil system. The operon structure of the kil genes is highly complex, with 5 different operons encoding for the kil genes. The operon with genes kilA to kilD contains 5 genes in total, including the polA gene. The initially proposed operon with 10 genes (Seef *et al*. 2021) was split in 3 by DiscOperon, with the first one harbouring the glyQ and glyS genes and 2 genes of unknown function, and one buffer gene containing an FHA-domain protein, all of which are encoded on the same strand. This highlights the importance that gene distance and neighbourhood is not sufficient to identify operons and that data on co-expression can be instrumental in distinguishing operonic from non-operonic genes. The second operon is the largest operon detected in the M. xanthus genome and contains 20 genes. Among those are kilB, kilG and kilH, as well as the minor pilin genes for Tip 4, kilK_4_, kilL_4_, kilN_4_ and kilM_4_. The other minor pilins encoding Tips 1, 2 and 3 are harboured on one operon each, whereby the single kilO gene, which is required for Tip3, is encoded together with kilM_3_, kilN_3_ and kilL_3_. See **Supplementary Table S3** for gene accession numbers and genomic positions in operons.

The **trp** operon encodes genes required for synthesizing the amino acid L-tryptophane (Trp). The enzymes involved in the biosynthesis of this amino acid are highly conserved, their genomic organisation, however, varies greatly between different bacterial species, from complete operons to complete dispersion of enzymes (reviewed in (Merino, Jensen, and Yanofsky 2008)). DiscOperon reconstructed the **trp** operon in *M. xanthus* as three transcriptional units. The first one contained the genes aroF, together with trpA to trpD and trpF, while the two other units each contained a copy of trpG and trpE genes **(Figure 3B)**.

Finally, we analysed the **kil** system that contains the genes in *M.xanthus* that are involved in predation (**Figure 3C**). The kil system contains a tad like pilus composed of a major pilin kilP, a prepilin peptidase kilA, a secreting kilC, an ATPase kilF, multiples set of minor pilins composing the Tips 1 to 4 (kilL_1-4_, kilM_1-4_, kilN_1-4_, kilK and kilO), the inner membrane platform composed of kilH and kilG, the outer membrane pilus assembly protein kilB, and kilD, a protein recruited at the contact site (Herrou *et al*. 2025). Most of the genes of the kil system are encoded in two large genes clusters MXAN_RS15020 - MXAN_RS15075 and MXAN_RS22530 - MXAN_RS22635 (described previously in (Seef *et al*. 2021)). Three additional clusters encode the first three sets of minor pilins, with kilO being two genes upstream of kilL_3_. kilP is located in another region of the genome. DiscOperon predicted the first cluster (MXAN_RS15025 - MXAN_RS15070) as two transcriptional units. The first includes MXAN_RS15025 - MXAN_RS15040, which encodes the genes glyQ and glyS, both required for glycine-tRNA ligation, as well as 2 unrelated genes of unknown function. This cluster is therefore not part of the kil system. MXAN_RS15045 is predicted as a monocistronic transcriptional unit and encodes an FHA-domain protein. Finally, kilA, kilC, kilF and kilD are grouped together in the same operon with MXAN_RS15070, which encodes the DNA polymerase polA. The second cluster (MXAN_RS22530 - MXAN_RS22635) was recovered in the largest predicted operon, which includes kilH, kilG, kilB/cpaB, kilL_4_, kilM_4_ and kilN_4_, as well as 4 hypothetical proteins. The remaining three minor pilins were each reconstructed as single operons together with a small number of other genes. KilO, which is only mandatory for predation if the third set of minor pilins is used (Tip3), is encoded in the same operon as the kilL_3_, kilM_3_ and kilN_3_ minor pilin set.

We looked at the kil system in more details concerning its syntenic conservation. A detailed and high-quality analysis was already done in Herrou et al. (Herrou *et al*. 2025), focusing on the conservation and operon structure of the kil system in a selection of predatory bacteria. To complement this, we performed fast.genomics analysis (Price and Arkin 2024) of the 5 kil operons (**Suppl. Figures S2-S6**). There is a strong association of kilA and kilC with kilF throughout evolution, while kilF is more often not found in the same operon (**Suppl. Figure S2 A**). The operon harbouring kilB, kilG, kilH and kilK, together with the 4^th^ tip is highly complex. Closely conserved seems the kilB-kilG-kilH unit, while kilK and the 4^th^ tip are not found with higher genome divergence (**Suppl. Figure S2 B**). For the other 3 tips, much less homologous genes were found with fast.genomics, except for the third tip. They seem however to be in close neighbourhood in nearly all identified species (**Suppl. Figure S2 C-E**).

In conclusion, using DiscOperon, we annotated the operon blocks contained in *M. xanthus* and could demonstrate that it can faithfully reconstruct known operon structures in this bacterium. All operons predicted in *M. xanthus* are available via download from Zenodo (https://doi.org/10.5281/zenodo.21976167), are available upon request from the authors and have been submitted to ODB.

## DISCUSSION

In this study, we developed DiscOperon, an open-source operon prediction tool that combines transcriptomic data and comparative genomics to reconstruct bacterial operons. Across multiple bacterial species, DiscOperon achieved accurate operon prediction and was able to faithfully predict operons of bacteria from diverse phyla.

We chose to combine genomic organisation, transcriptomic evidence and comparative genomics to faithfully reconstruct the operon structure in bacterial genomes, which brings several advantages. Gene co-expression information provides direct evidence, if neighbouring genes are transcribed together across different experimental conditions. Evolutionary conservation, on the other hand, reflects functional relationships of gene neighbours that are maintained across species. By combining these complementary features, DiscOperon is able to distinguish true operonic gene pairs from adjacent genes that are only co-expressed by chance. The integration of comparative genomics during the operon reconstruction step furthermore enables our tool to not only consider directly adjacent genes, but to also retrieve operon information despite gene insertions, deletions or local rearrangements. We consider this an important feature, as it better accounts for the complex evolutionary processes taking place at the level of genome organisation in bacteria.

We applied DiscOperon on the genome of *M. xanthus* to annotate its operon structure. Based on our analysis of well-characterized operons, our tool could faithfully identify *M. xanthus* operon structures: despite its large and highly duplicated genome, we were able to correctly retrieve different operons involved in diverse biological processes, such as the trp and frz operons or the different gene clusters of the kil system.

This confirms DiscOperon’s ability to predict operons in phylogenetically diverse bacteria and in less well characterized bacteria with a complex evolutionary history.

DiscOperon achieves similar, or better results than currently available methods, such as OperonFinder, OpDetect or Rockhopper, when comparing accuracy of detecting gene pairs. We want to point out that DiscOperon returns directly the list of operons and their associated genes, and is an easy, light-weight program applicable to any bacterial species. Furthermore, given that the problem of operon detection in genomes is rather old, it is noteworthy that only few tools that were developed and published over the decades are still functional. We chose those three for comparison, as they were the ones which were in our hands the tools that could still be used for operon prediction without major issues.

Still, DiscOperon has several limitations, the most important ones we discuss below. As with any transcriptomics-based approach, operon prediction quality depends on the diversity and quality and the number of different transcriptomic data used: using only few datasets, or data retrieved under similar experimental conditions will most likely result in less reliable results. Furthermore, operons that are inactive in the different conditions used, or which contain genes that are rarely active, may be difficult to detect. We therefore recommend using a sufficiently large set of data from different experimental conditions. Furthermore, while comparative genomics makes operon prediction more robust, it also makes it more difficult to predict operons for genes with few homologs in the chosen reference genomes, or for bacteria with long evolutionary distances to available sequenced genomes. We also have not tested DiscOperon on species with only incomplete genome assemblies.

## DATA AND CODE AVAILABILITY

We designed DiscOperon as a lightweight command line tool requiring minimum programming skills to be used. It can be run on a regular computer without requirement of GPU or high-performance computing power. The code is freely available at: https://gitlab.com/habermann_lab/discoperon. All files used as input are publicly available: Genomes and protein sequences were extracted from the Refseq database and RNA-seq data from the sequence read archive (SRA), both from NCBI. The GEO accession numbers of the different RNA-seq experiments used in this study can be found in **Supplementary Table S2**.

## ACKNOWLEDGEMENTS

We thank the Computational Biology team at IBDM for helpful discussions.

## FUNDING

TB was financed by a MENTR PhD fellowship from Aix-Marseille University and Fondation de la Recherche Medicale (FRM) -grant FDT202504020470. This work was supported by the IBDM (UMR7288), the Centre Nationale de la Recherche (CNRS) and by Aix-Marseille University (AMU).

## AUTHOR CONTRIBUTIONS

This study was conceived by TB and BHH. TB implemented code, did all computations and analysed the data, wrote the manuscript and prepared a first draft of all figures. BHH supervised the study, helped with data analysis and interpretation, wrote the manuscript and finalized figures for publication.

## Notes

https://doi.org/10.5281/zenodo.21976167

https://gitlab.com/habermann_lab/discoperon

### Competing Interest Statement

The authors have declared no competing interest.

