## Supplementary Figures for "Predicting the operon structure of the *Mycococcus xanthus* genome using the novel software DiscOperon"

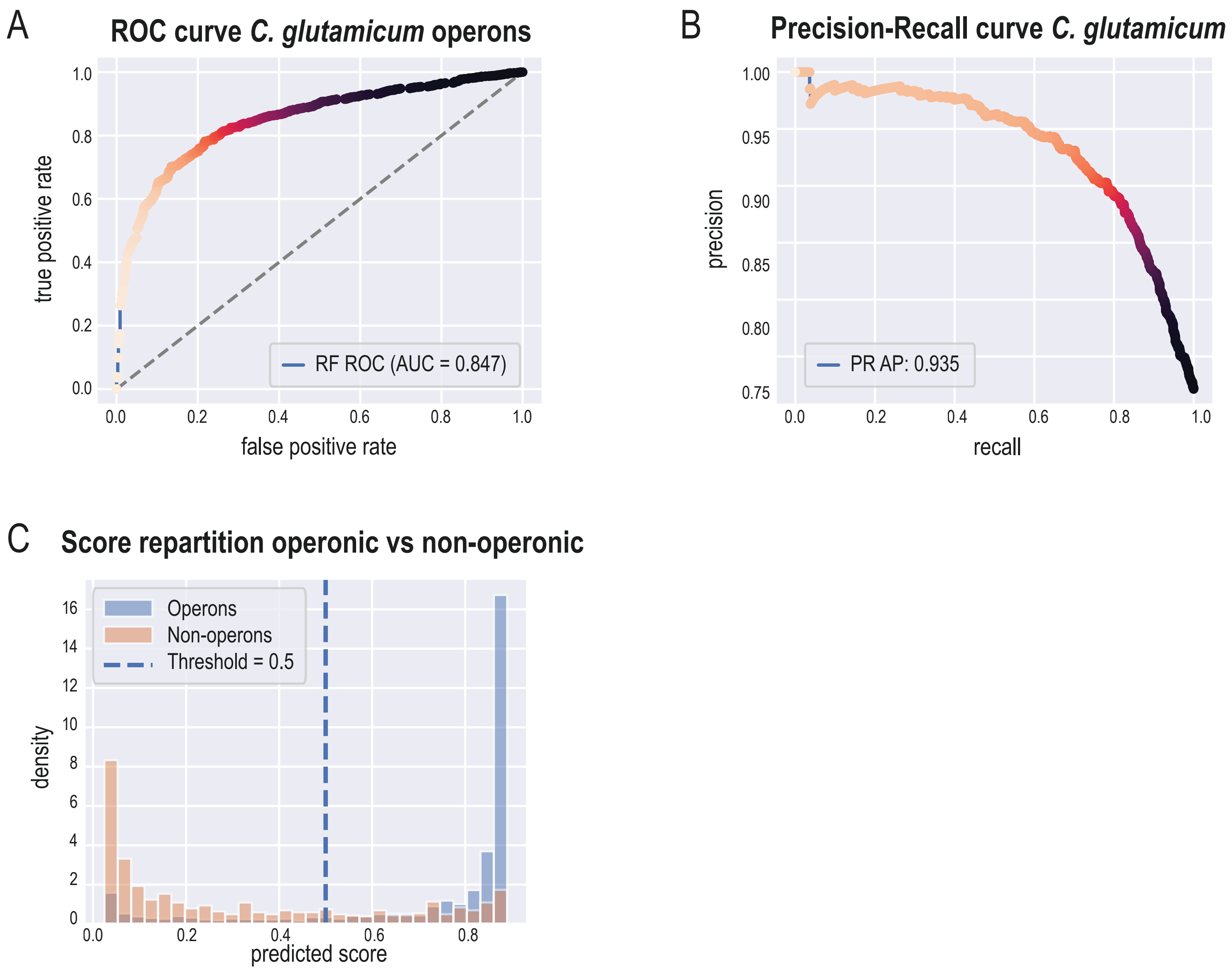


**Figure 2: Performance of DiscOperon in correctly predicting operons in *C. glutamicum***. (A) ROC curve relating true-positive and false-positive prediction rates, and (B) Precision-Recall curve of operons predicted for *C. glutamicum*. The ROC area under the curve (AUC) was 0.847, the average precision (AP) was 0.935. (C) Score repartition of operons vs non-operons. Operons (blue bars) are clearly overrepresented at prediction scores > 0.5, while non-operons are clearly overrepresented at lower scores, demonstrating DiscOperon's ability to correctly distinguish operonic vs non-operonic gene pairs.

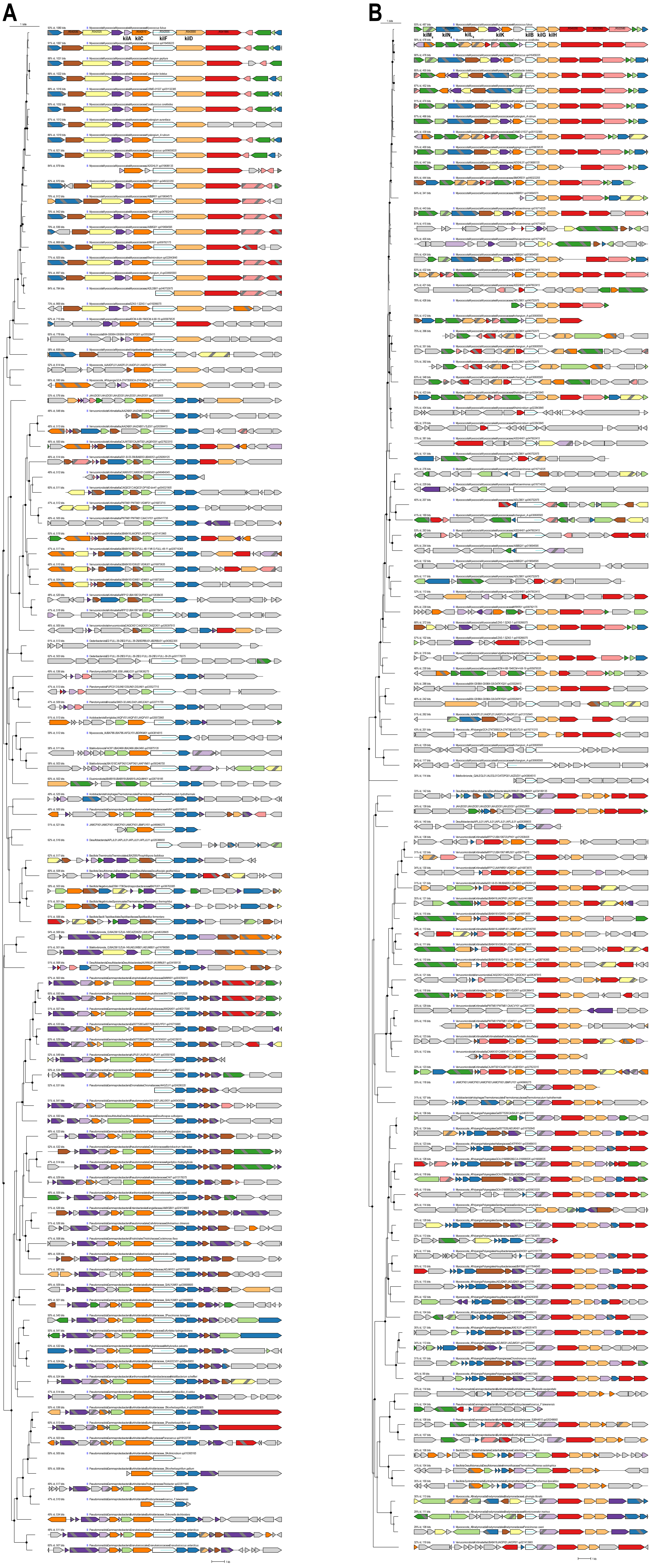


**Supplementary Figure S2 A and B: Synteny conservation of the two major kil system operons in *M. xanthus*. (A)** Operon structure conservation of the kil-gene cluster I. kilF was used as input for a fast.genomics search (https://fast.genomics.lbl.gov/cgi/search.cgi).
The kilC and kilA genes have stronger association with kilF than kilD.

**(B)** Operon structure conservation of the kil-gene cluster II. kilB/cpaB was used as input for a fast.genomics search. The inevlution. structure of the kilB, kilG and kilH cluster seems highly conserved in evolution. The white arrow with the blue line is the query gene, identical colors indicate homologous genes.

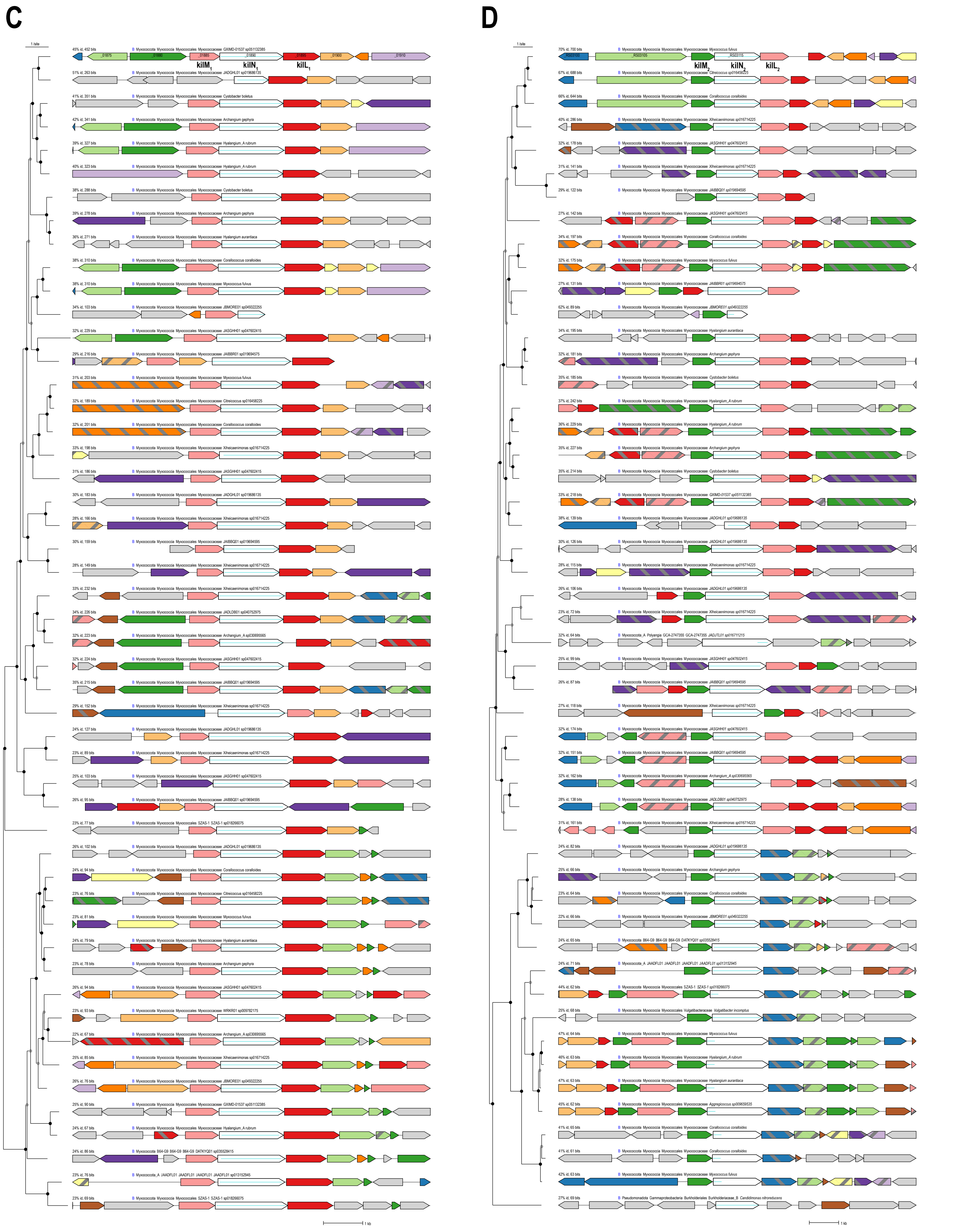


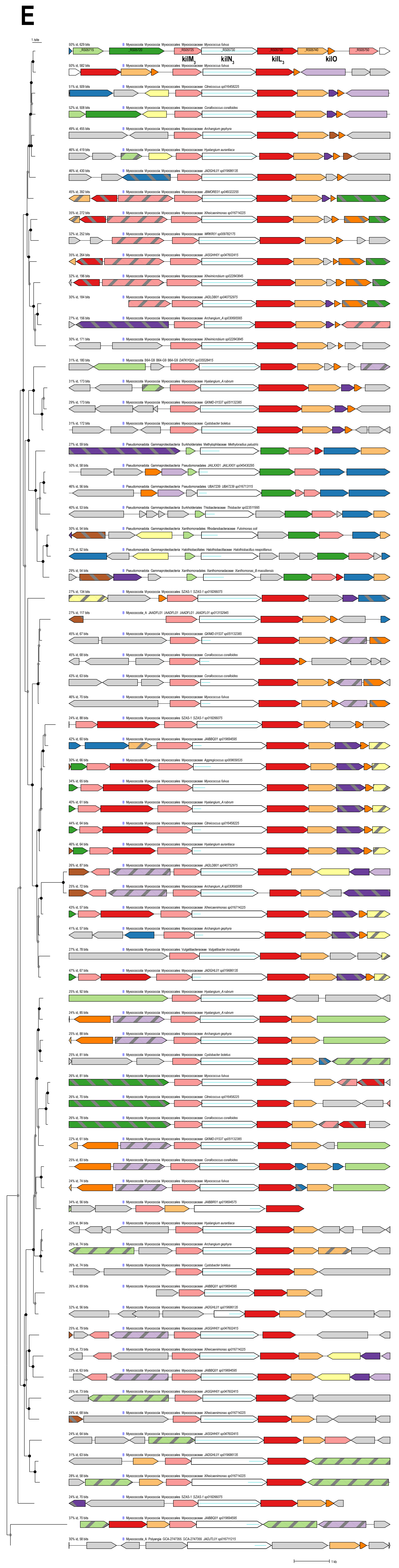


**Figure S2 C, D and E: synteny conservation of the minor pilin gene clusters. (C)** Synteny consevation of the Tip1 genes kilL1, kilM1 and kilN1, as determined by fast.genomics. The three genes cluster well within the Mycococcota. **(D)** Synteny conservation of the Tip2 genes kilL$_2$, kilM$_2$ and kilN$_2$. More divergence is seen here and only kilM$_2$ and kilN$_2$ are coupled, kilL$_2$ is either lost from the neighborhood or has been relocated within the neighborhood. **(E)** Synteny conservation of the Tip3 genes kilL3, kilM3, kilL3 and kilO. More conserved blocks could be found with these genes. Relatively high conservation is found of the neighborhood, the kilO genes is not always found. The kilN1, kilN2 and kilN3 genes were used for fast.genomics searches and are indicated as white arrows with a blue line. Same color in the individual figures indicate conserved genes.
